# Emerging parvovirus associated with an outbreak in Dutch pig farms also detected in pigs and wildlife in Denmark

**DOI:** 10.64898/2026.06.24.734170

**Authors:** Marta Canuti, Frederikke Juncher Høeg, Anne Sofie Vedsted Hammer, Tim Kåre Jensen, Michelle Lauge Quaade, Pia Ryt-Hansen, Aida Droce, Charlotte Mark Salomonsen, Simon Smed Sørensen, Lars Erik Larsen

## Abstract

A parvovirus recently associated with an outbreak in Dutch pigs was found in Denmark in symptomatic pigs and in fox (*Vulpes vulpes*) feces and spleens. Pig viruses were more closely related to each other than to viruses found in the respective local wildlife, suggesting a link between the farm outbreaks.

---

Recently, a parvovirus similar to the one discovered in fecal samples from Dutch red foxes (fox parvovirus, species *Protoparvovirus carnivoran4*) (1) was responsible for a multi-farm outbreak associated with exophthalmos, erythema, and other nonspecific clinical signs in piglets from >80 conventional herds in the Netherlands (2).

In June 2026, 1.5-2 % of piglets and weaners in a Danish herd with 920 sows experienced similar clinical signs, including exophthalmos, strabismus, erythema, and alopecia (Figure 1). The herd annually tested free of *Mycoplasma hyopneumoniae, Actinobacillus pleuropneumoniae* 2, 6, and 12, porcine reproductive respiratory syndrome virus 1 and 2, *Brachyspira hyodysenteriae, Pasteurella multocida, Sarcoptes Scabiei Suis*, and *Haematopinus suis*. Nursery pigs were treated with antibiotics without any effect. Some sows experienced fever probably due to swine influenza, detected in the herd. Six piglets (aged 1-2 weeks) were submitted for necropsy. Macroscopic examination revealed normal body conditions with no alopecia or erythema. Mild exophthalmos, eyelid edema, and mild ocular discharge were seen in four animals. All showed rhinitis (mucus in the nasal cavity); one piglet had consolidation of a lung lobe (*lobus medius*, ~ 7% of the lung tissue) and serous content in one bulla tympanica. Two piglets had little stomach content, four had moderate amounts of milk. Two piglets showed increased synovial fluid in several joints, and streptococci were cultured from one elbow joint. Liver, spleen, and kidneys were swollen, possibly because of euthanasia with pentobarbital. Livers and kidneys from all six pigs tested positive for the emerging parvovirus (DNA isolations and semi-nested PCR as in (3) with primers: Fox_Parvo_F1:ACTGTGGTTCGCATAGGCTG, Fox_Parvo_R4:CTTGGATAGTAGACCAGGTTG; Fox_Parvo_R1:GTTCAGCAGTTGGCTTGGTG).

**Figure 1.**
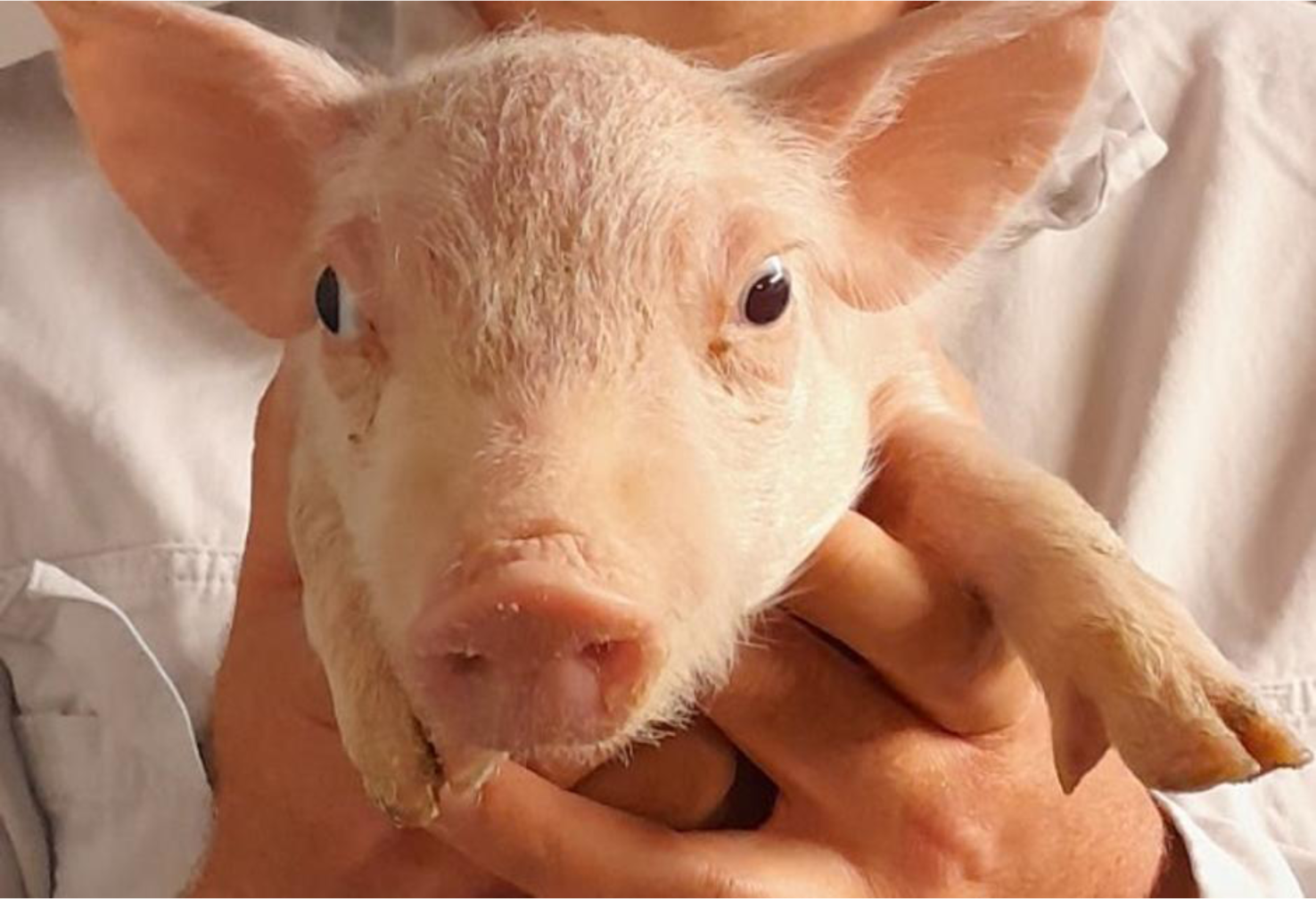
Typical clinical appearance of the affected pigs showing clear exophthalmos and eyelid oedema.

DNA previously isolated from Danish wild animals (3) was also screened. The virus was detected in one of two fecal samples from red foxes (*Vulpes vulpes*), confirming the results by Bodewes and colleagues in the Netherlands (1) and by Lojkić et al., who found this virus in fecal samples from Croatian foxes (4). Additionally, the virus was detected in two out of 80 spleen samples from Danish red foxes (2.5%) but in none of the spleens from raccoon dogs (*Nyctereutes procyonoides*, N=3), Eurasian otters (*Lutra lutra*, N=10), beech martens (*Martes foina*, N=6), European badgers (*Meles meles*, N=5), pine martens (*Martes martes*, N=2), and polecat (*Mustela putorius*, N=1). The two positive foxes did not show any pathological signs; one was amdoparvovirus-positive (3).

The full viral coding genome from the fox fecal sample (XFM39) was obtained through metagenomics (18,876 reads, average coverage: 558X) (5), while those of the viruses identified in a fox spleen (FFM99) and two pig livers (FPP1 and FPP4) were obtained through overlapping PCRs and Sanger sequencing (primers available upon request). From the other samples, partial sequences were obtained by sequencing the screening PCR products: the virus from the third fox was identical to XFM39, and those from pigs were 100% identical to FPP1/FPP4. Maximum likelihood phylogenetic trees (6) showed that the virus from the Dutch outbreak was the closest relative to the ones from the Danish pig farm (99.9% genomic identity, two non-synonymous mutations in the non-structural protein, and none in the capsid protein), and both were included in a clade encompassing also Dutch and Croatian fox viruses (Figure 2). Differently, the genomes of the viruses from Danish foxes were <95% identical to all other viruses and formed a distinguished clade.

**Figure 2.**
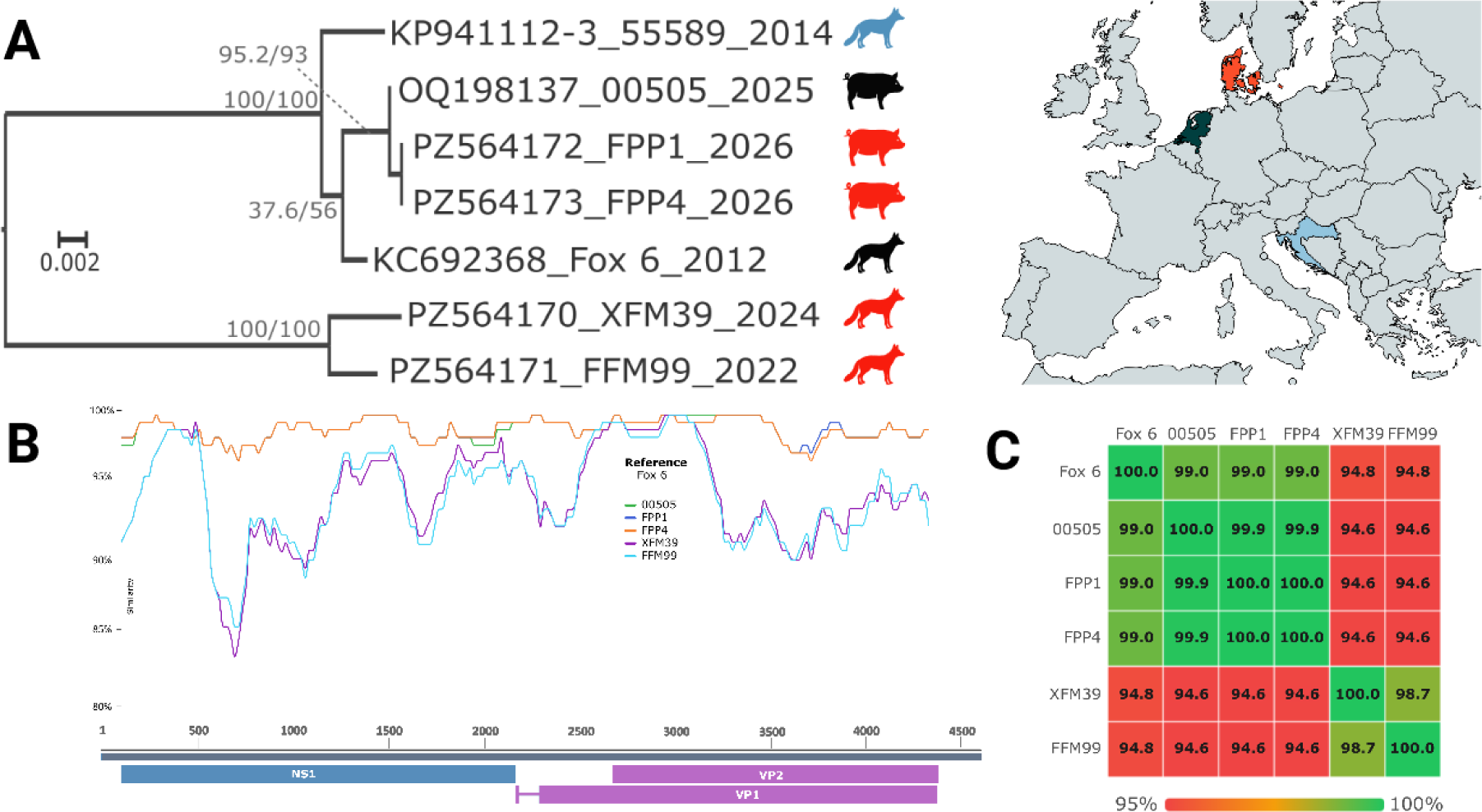
Sequence and phylogenetic analyses of viruses within *Protoparvovirus carnivoran4*. A) Phylogenetic analysis of concatenated (1152 nt) non-structural (NS1) and structural (VP2) genomic fragments of viruses identified in foxes and pigs, as indicated by the animal logos, which were colored following the country of origin of the virus according to the map on the right. Each virus sequence’s name is labelled by the GenBank accession number, the strain name, and the year of detection. The full genome was not used as partial sequences are available for virus 55589, but analyses with full genomes and genes gave consistent results. The outcomes of the Shimodaira–Hasegawa approximate likelihood-ratio test and ultrafast bootstrapping (1000 replicates) are indicated at the nodes. The tree, midpoint rooted, was built with IQTree3 according to the TPM2u+F model, identified as the best fitting for genetic distance estimation (modelfinder function). B) Similarity comparisons across pairs of complete genomes of viruses from pigs and foxes compared to the query sequence Fox 6. The y-axis depicts pairwise identity (Window: 200, Step: 20) across the genome, whose positions are reported on the x-axis, illustrated at the bottom. Colored rectangles represent predicted open reading frames after splicing. C) Pairwise sequence identity (percentage) between the available full genomes, where values are color-coded according to the legend at the bottom. All analyses were performed with Stranded (VerCan ApS), the map was created with MapChart ©, and the final image was created in BioRender (Canuti, M. (2026) https://BioRender.com/beuq540).

Our study confirms that this virus is possibly a porcine pathogen and reports for the first time its detection in Denmark. The observed clinical signs resembled those seen in several Dutch herds, further indicating that this is a disease syndrome not described before. Further controlled studies are needed to establish viral prevalence and its association with disease.

The detection of this virus in the fox spleens strongly indicates that it can infect and replicate in this species, although its pathogenic role in wildlife remains unknown. Viruses from Dutch and Danish pig farms were more identical to each other than to viruses found in the respective local wildlife. This indicates an epidemiological link between the farm cases and supports a single introduction into pigs with spread into multiple countries. While the detection of this virus in foxes predates the recognition of the associated pig syndrome, it is not possible to definitely conclude whether foxes and pigs were infected by the same source, the virus originated from wild animals and was introduced into farms, or vice versa, as past farm outbreaks could have been missed. However, the higher viral diversity in foxes might suggest a fox origin. As parvoviruses are extremely stable (7), we can hypothesize an environmental-related transmission between wild and farmed animals (e.g., through contaminated material introduced into farms or through exposure to infected farm-originating carcasses). Further studies are required to assess the origin and spread of this virus and clarify its ecology, including identifying its maintenance and spillover host(s) and host range.

## Acknowledgments

We thank Joost TP Verhoeven for the implementation of the metagenomic pipeline V2D and Nina Dam Grønnegaard, Hue Thi Thanh Tran, Anna Cecilie Boldt Eiersted, and Sally Xin Hildebrandt for technical assistance.

## About the Author

Marta Canuti is an Assistant Professor at the University of Copenhagen. Her primary research interests are virus discovery and emergence, and her studies focus on virus epidemiology, ecology, and evolution. She primarily investigates virus transmission dynamics and cross-species transmission in wildlife and at the interface between wildlife and domesticated animals.

